# An atomistic model of the coronavirus replication-transcription complex as a hexamer assembled around nsp15

**DOI:** 10.1101/2021.06.08.447516

**Authors:** Jason K. Perry, Todd C. Appleby, John P. Bilello, Joy Y. Feng, Uli Schmitz, Elizabeth A. Campbell

## Abstract

Using available cryo-EM and x-ray crystal structures of the nonstructural proteins that are responsible for SARS-CoV-2 viral RNA replication and transcription, we have constructed an atomistic model of how the proteins assemble into a functioning superstructure. Our principal finding is that the complex is hexameric, centered around nsp15. The nsp15 hexamer is capped on two faces by trimers of nsp14/nsp16/(nsp10)_2_, where nsp14 is seen to undergo a large conformational change between its two domains. This conformational change facilitates binding of six nsp12/nsp7/(nsp8)_2_ polymerase subunits to the complex. To this, six subunits of nsp13 are arranged around the superstructure, but not evenly distributed. Two of the six polymerase subunits are each proposed to carry dimers of nsp13, while two others are proposed to carry monomers. The polymerase subunits that coordinate nsp13 dimers also bind the nucleocapsid, which positions the 5’-UTR TRS-L RNA over the polymerase active site, a state distinguishing transcription from replication. Analyzing the path of the viral RNA indicates the dsRNA that exits the polymerase passes over the nsp14 exonuclease and nsp15 endonuclease sites before being unwound by a convergence of zinc fingers from nsp10 and nsp14. The template strand is then directed away from the complex, while the nascent strand is directed to the sites responsible for mRNA capping (the nsp12 NiRAN and the nsp14 and nsp16 methyltransferases). The model presents a cohesive picture of the multiple functions of the coronavirus replication-transcription complex and addresses fundamental questions related to proofreading, template switching, mRNA capping and the role of the endonuclease. It provides a platform to guide biochemical and structural research to address the stoichiometric and spatial configuration of the replication-transcription complex.

**Author Summary:** The replication of the coronavirus genome and the synthesis of subgenomic mRNA is a complex process involving multiple viral proteins. Despite a fairly complete structural picture of the individual proteins that are believed to coalesce into a larger replication-transcription complex, there is no clear model of how these proteins interact. Here we present the first detailed atomistic model of a complete replication-transcription complex for SARS-CoV-2, made up of the non-structural proteins nsp7-nsp16, as well as the nucleocapsid. Forming a large, hexameric superstructure centered around nsp15, the model provides new perspective on the function of its individual components, including the exonuclease, the endonuclease, the NiRAN site, the helicase, the multiple zinc fingers, and the nucleocapsid. It offers a cohesive view of replication, proofreading, template switching and mRNA capping, which should serve as a guide for future experimental exploration.

## Introduction

Coronaviruses, including the SARS-CoV-2, pathogen responsible for COVID-19, constitute a family of positive sense, single stranded RNA viruses. After cellular infection, these viruses harness both viral proteins and host machinery to reproduce the viral genome and assemble new viral particles that are released to infect new hosts.[1] The replication and transcription of the coronavirus genome is facilitated by a complex of nonstructural viral proteins (nsp’s) encoded by the *ORF1ab* gene (Figure 1). The *ORF1a* gene translates a polyprotein which spans nsp1 to nsp10. This polyprotein is cleaved into discrete proteins by two encoded proteases, nsp3 and nsp5. After a frameshift, a longer *ORF1ab* polyprotein may be produced, which includes the additional proteins nsp12 to nsp16. This set of proteins, in addition to the nucleocapsid (N) which packages the viral RNA, perform the primary functions of RNA synthesis, proofreading, mRNA capping, and strand separation in both full genome replication mode and sub-genomic transcription mode.

**Figure 1.**
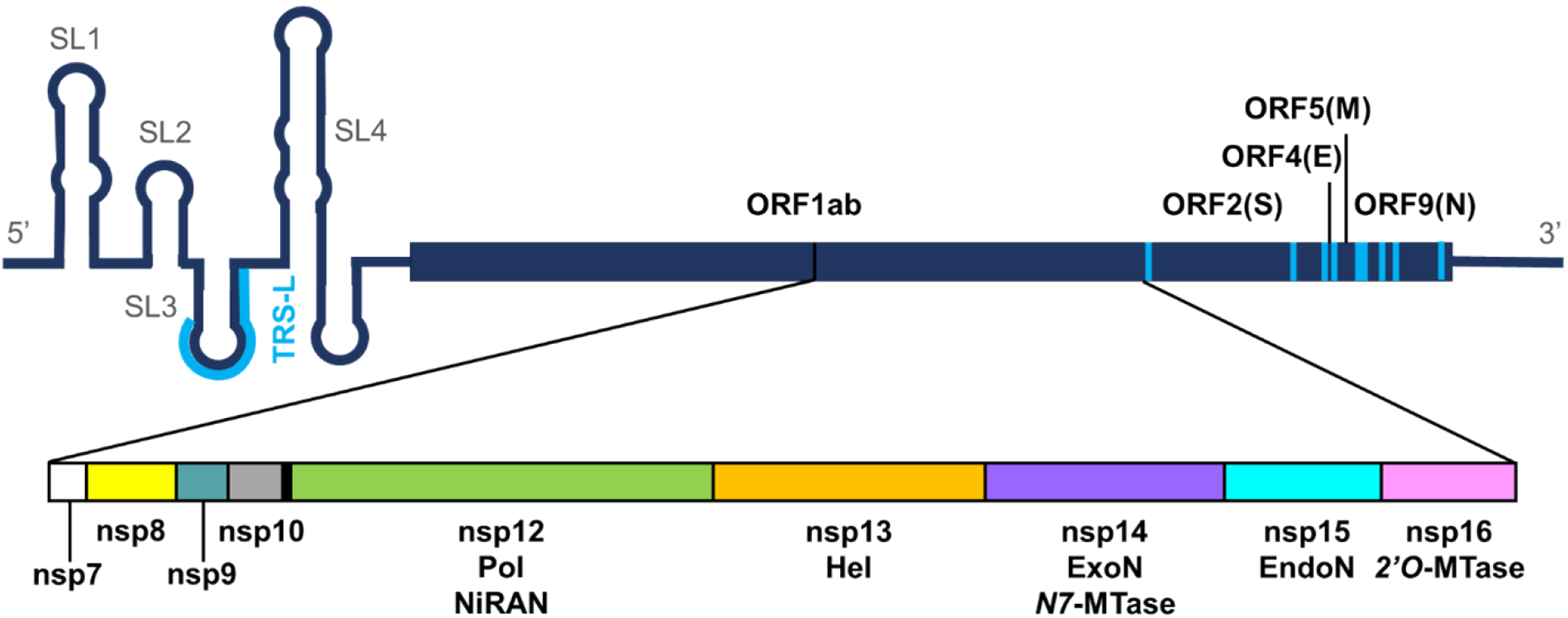
Organization of the SARS-CoV-2 genome. The genome is divided into multiple open reading frames (ORF’s), with ORF1ab containing the non-structural proteins (nsp’s) required for RNA replication (nsp7 – nsp16). The structured 5’-UTR leader contains a transcription regulatory sequence (TRS-L) which is repeated throughout the genome, each instance preceding an ORF (indicated in blue). During negative-strand synthesis, shorter transcripts may be generated when the template switches from one of the TRS locations in the body of the genome to the TRS-L location in the 5’-UTR, effectively skipping over everything in between.

The majority of these proteins have been well characterized. Nsp12 contains the polymerase (Pol) active site, responsible for RNA synthesis.[2] It also has a second enzymatic site referred to as the NiRAN (nidovirus RdRp-associated nucleotidyltransferase), which is unique to viruses in the *Nidovirales* order.[3] The role of this site remains ill-defined, but it has been shown to nucleotidylate proteins such as nsp9,[4] and it has been proposed to play the critical role of guanylyltransferase (GTase) in mRNA capping.[5] Nsp14 also has two enzymatic activities.[6] Its N-terminal domain (NTD) is an exonuclease (ExoN), which has been shown to be responsible for RNA proofreading during synthesis. Its C-terminal domain (CTD) is an *N7*-methyltransferase (MTase) involved in mRNA capping. Nsp16 is a *2’O*-MTase, also involved in mRNA capping.[7] Nsp13 is a helicase (Hel), capable of unwinding RNA, powered by a nucleoside triphosphatase (NTPase) site.[8] Nsp15 is an endonuclease (EndoN), which is believed to cleave the 5’ poly-U tail of the intermediate negative-strand.[9] And finally, various cofactors such as nsp7, nsp8, nsp9, nsp10 and the nucleocapsid (N) have been shown to be involved in replication and transcription as well.[10-13]

Unlike many other viral families, structures have been determined for all key proteins that are presumed to make up the coronavirus replication-transcription complex (RTC), several of which are shown in Figure S1. To date, there are x-ray crystal structures of coronavirus nsp13,[14, 15] heterodimeric nsp14/nsp10,[6, 16, 17] heterodimeric nsp16/nsp10,[11, 12, 18-21] hexameric nsp15,[22-27] dimeric nsp9,[28-30] and the N protein NTD bound to both dsRNA and the specific viral RNA oligo known as the transcription regulatory sequence (TRS), which is critical to the unusual template switching process that occurs during transcription.[31] Cryo-EM has been especially successful in illuminating the structure of the polymerase complex, made up of nsp12, nsp7, two subunits of nsp8, and up to two subunits of nsp13.[2, 5, 10, 32-36] In this complex, nsp7 and the two nsp8 subunits sit atop the nsp12 Pol active site, coordinating to the thumb and fingers domains. The long and flexible N-terminal (N-term) nsp8 helices extend out over the exiting dsRNA when the complex is captured in its replicating state. Coordinated to these nsp8 helices, two subunits of nsp13 sit above the polymerase complex, where one of them has been observed engaging the downstream RNA template overhang.

Yet despite this wealth of structural information, there is no clear picture of how the polymerase complex interacts with the remaining proteins to form the complete RTC, leaving major questions of viral RNA processing unanswered. Functionally, it is inferred that the nsp14 ExoN must have some interaction with nsp12 in order to gain access to the 3’ end of the nascent strand to carry out proofreading during RNA synthesis. The dsRNA that emerges from the polymerase must eventually be unwound. To produce transcripts for protein translation, the resulting 5’ end of the nascent positive-strand must be directed to the mRNA capping sites: presumably the NiRAN site of nsp12 and the two MTase sites of nsp14 and nsp16. The complicated process of template switching during transcription remains entirely enigmatic but appears to involve N recognizing the specific junctions where the switch occurs: the leader TRS (TRS-L) and body TRS (TRS-B) (Figure 1).[37]

In this work, we have endeavored through molecular modeling to determine how the many proteins identified above assemble into a functioning SARS-CoV-2 RTC. The key to unlocking this puzzle was the realization that nsp15 could provide the scaffolding around which the complex could be built. Nsp15 is perhaps the least studied of the enzymes encoded by *ORF1ab*. Its EndoN activity appears to be dispensable, as mutations which render the EndoN catalytically dead, still lead to viable, albeit less robust, viral replication.[38] However, those same studies showed nsp15 mutations outside of the EndoN active site have lethal consequences to the virus. Considering coimmunoprecipitation studies also showed that nsp15 strongly colocalizes with proteins of the RTC,[39] collectively, these data suggest nsp15 plays a structural role that is critical to the proper function of the RTC.

With this as the starting hypothesis, assembly of an atomistic model of the complex from the existing structures turned out to be relatively straightforward. The resulting RTC superstructure is a hexamer, with six subunits each of nsp12, nsp13, nsp14, nsp15 and nsp16. The model demonstrates how RNA makes its way from the Pol active site, across the ExoN and EndoN sites, separates into template and nascent strands, and directs the 5’ end of the nascent strand to the three mRNA capping sites. We’ve also identified a binding site for the TRS-L bound N protein that has implications for template switching during negative-strand synthesis.

Here we outline how the model was constructed and its overall architecture. We describe the implications on multiple functions associated with the RTC, including proofreading, mRNA capping, template switching, and negative-strand poly-U cleavage. We believe this work presents a cohesive picture of the complicated processes associated with coronavirus genome replication and transcription and offers a roadmap for further exploration.

## Results

### Proteins comprising the RTC

The model was assembled from existing structures of the individual SARS-CoV-2 components (Figure S1). This included the known SARS-CoV-2 polymerase complex nsp12/nsp7/(nsp8)_2_/(nsp13)_2_/dsRNA (PDB:6XEZ)[32] (Figure S1a), an homology model of nsp14/nsp10 based on a SARS-CoV x-ray structure (PDB:5NFY, chains A/M)[6] (Figure S1b), nsp16/nsp10 (PDB:6WVN) [21] (Figure S1c), hexameric nsp15 (PDB:6X1B)[25] (Figure S1d), dimeric nsp9 (PDB:6W4B),[40] and the NTD of the N protein bound to the 10-nucleotide (nt) TRS-L oligo (PDB:7ACT)[31] (Figure S1e). Following previous precedent, we refer to the two subunits of nsp8 in the polymerase complex as nsp8.1 and nsp8.2, and the two subunits of nsp13 as nsp13.1 and nsp13.2.[32] Similarly, as there are two sources for nsp10 (associated with nsp14 and nsp16), we refer to them as nsp10.14 and nsp10.16, as necessary for clarity. Finally, as the hexameric nature of the complex turned out to be critical, we adopt a notation based on the nsp15 structure to simplify the discussion. This structure can be viewed as a dimer of trimers (see Figure S1d), presenting two trigonal faces: face A and face B. We refer to the subunits of nsp15 as A1, A2, A3, B1, B2, and B3. This notation will subsequently be used to identify associated subunits of the superstructure.

### Initial protein-protein docking

We undertook a systematic approach to dock various components of the complex in a binary fashion. The outcome of most of these exercises was not particularly fruitful, providing no new insight into the interactions between the polymerase complex and nsp14/nsp10, monomeric nsp15 or nsp16/nsp10. Most of the resulting docking poses for these key proteins suggested more of a preference for interaction with the dsRNA exiting the polymerase than interactions with the proteins of the polymerase complex itself. Similarly, little insight into the interactions within the set of nsp14/nsp10, nsp15, and nsp16/nsp10 could be discerned.

However, two exceptions emerged. The nsp9 dimer, a putative RNA binding protein, demonstrated a clear preference for binding to the 2A domain of nsp13 (Figure S2). This proved to be a consistent finding, as we followed up with docking of the nsp9 dimer to several available structures of nsp13, capturing a variety of conformations of this protein. Notably, the position of nsp9 on nsp13 is such that it is ideally situated to interact with the 5’ end of the ssRNA as it exits the helicase RNA binding groove.

Additionally, we found that the NTD of the N protein, as bound to a 10-nt oligo corresponding to the TRS-L of the SARS-CoV-2 genome, binds robustly between the two nsp13 subunits of the polymerase complex (Figures 2a and 2b), as all thirty of the returned binding modes were variations on this same interaction. This positions the TRS-L above the Pol active site, having implications for template switching during transcription. To emphasize this point, the top scoring docking pose orients the TRS-L RNA parallel to the polymerizing template strand, with the 5’ end on the entrance side of the Pol active site and the 3’ end on the exit side. The C-terminus (C-term) of this domain, as defined by residue S180, is also well exposed to solvent on the exit side of the polymerase, implying that full-length N protein could bind unimpeded. Interestingly, we found that structures of the N protein that are either apo or dsRNA binding do not dock to this site. Only the structure with the co-crystalized TRS-L segment binds here, which is likely due to its more spherical shape.

**Figure 2.**
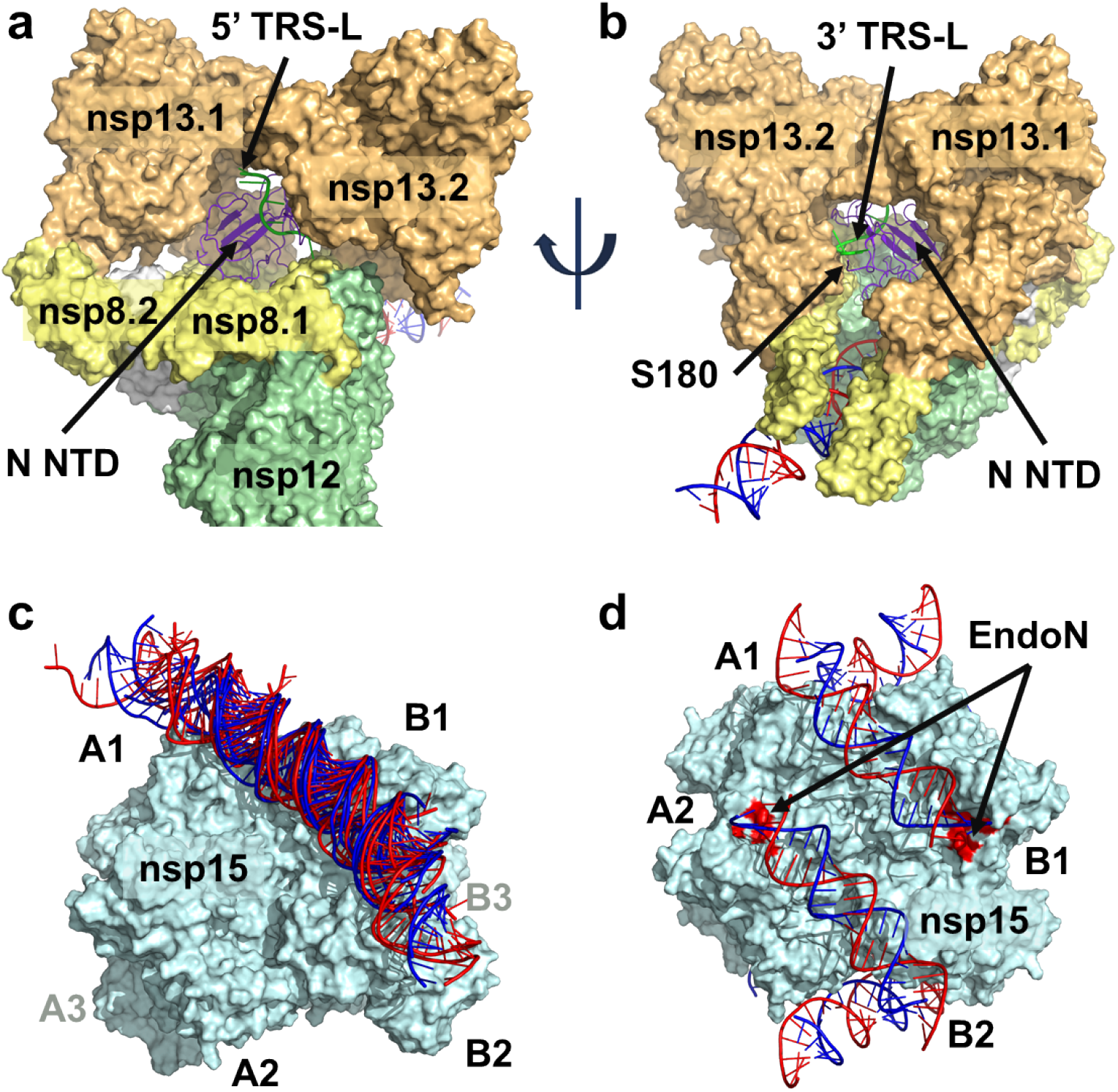
Significant findings from protein-protein docking. (a,b) The N NTD bound to a TRS 10 nt oligo (PDB: 7ACT) docks into the void between the two nsp13 subunits of the polymerase complex. The TRS oligo is positioned over the polymerase active site, parallel to the template, with its 5’ end exposed on the entrance side of the polymerase and its 3’ end exposed on the exit side. The C-term S180 residue of the N NTD is also exposed on the exit side of the polymerase, indicating full length N could bind to the complex unobstructed. (c) Multiple examples of dsRNA docked across the nsp15 hexamer, spanning subunit A1 to B1/B2. (d) Six dsRNA double helices can be symmetrically arranged around the hexamer: three directed from A → B, and three directed in an antiparallel fashion from B → A. Each dsRNA passes over an EndoN active site, colored in red.

### Nsp15-dsRNA interaction

Having completed this initial survey of potential binary protein-protein interactions, we considered the possibility that nsp15 retained its hexameric form in the RTC. Docking of the nsp15 hexamer to the polymerase complex did not identify any direct interaction between the proteins, but instead revealed a specific interaction between nsp15 and dsRNA that was common to all of the top poses. We followed up with docking of isolated dsRNA to the nsp15 hexamer and again found this particular binding mode was dominant (Figure 2c). The RNA runs from one face of the nsp15 hexamer to the other and is in contact with three subunits (A1, B1 and B2). As shown in Figure S3, it has direct interactions with at least eight basic residues (K110, R135 and K149 from subunit A1, K316, K319, K334 and K344 from B1, and K12 from B2) and passes over one of the EndoN active sites (on subunit B1). An appealing aspect of this finding is that we can arrange six dsRNA’s around the hexamer without interference (Figure 2d): three run from face A to face B and three run in the antiparallel direction from face B to face A. Each passes over a different EndoN site, suggesting to us that nsp15 could act as the core of the RTC, facilitating multiple replication cycles and directing RNA from the polymerase of one subunit to distal subunits that perform capping.

### Nsp15-nsp14/nsp10 interaction

An initial hypothesis we had with respect to proofreading was that the nsp14 ExoN interacted with the dsRNA as it exited the polymerase. We arrived at this conclusion based on an analysis of ExoN locations relative to the polymerase active site in various DNA polymerases. Notably, bacterial DNA polymerase I (e.g. the *E. coli* Klenow fragment)[41] and mammalian DNA polymerase γ[42] have the ExoN site located below the exiting dsDNA. Other potential locations of the ExoN domain, such as that seen in mammalian DNA polymerase δ,[43] are precluded by the known positions of nsp7, nsp8 and nsp13. Bolstering this hypothesis, the coronavirus ExoN has been shown to have a preference for dsRNA or hairpin ssRNA substrates over unstructured ssRNA.[11] Furthermore, a co-crystal structure of the similar Lassa ExoN provided a good example for how the nsp14 ExoN could interact with dsRNA.[44]

With this in mind, we considered that nsp14 could interact with the dsRNA as it is bound to the nsp15 hexamer. However, docking of nsp14 to this complex did not produce any meaningful poses. Manual exploration suggested a potential site of interaction where the ExoN was located under the dsRNA on the face opposite to the EndoN site (following the above example, with the EndoN site on B1, the ExoN could be positioned to interact with the RNA on A1). Intriguingly, this location allowed us to simultaneously position the zinc fingers of nsp10 over an antiparallel dsRNA just after it passes over the EndoN site (on A2). As attractive as this location was, the issue with this binding pose was that the MTase domain was not ideally situated with respect to nsp15. A similar situation arose when attempting to find an interaction between the ExoN and the exiting dsRNA of the polymerase complex. A location that functionally made sense for the ExoN, was not ideally positioned with respect to the MTase domain.

A sizable limitation of the protein-protein docking method employed is that it assumes the proteins have minimal flexibility and does not sample alternative conformations. The available structures of SARS-CoV nsp14 suggest at least some limited flexibility between the two domains exists.[6, 16] As they are linked by a 14-residue loop (residues 286-299), we considered the possibility that there may be greater flexibility between these two domains, an hypothesis supported by a recently published molecular dynamics (MD) study of SARS-CoV-2 nsp14[17] and made more likely in the context of protein-protein perturbations. Thus, given our assessment that the ExoN could interact with the dsRNA on both the polymerase complex and the nsp15 hexamer, and that it could potentially adopt a range of conformations with respect to the MTase, we decided to handle the two domains separately, first optimizing the position of the ExoN (residues 1-285) and nsp10 on nsp15 and then docking the MTase (residues 300-526).

We optimized the structure of ExoN/nsp10 interacting with nsp15 and dsRNA as described above (Figure 3a) and arranged six subunits symmetrically around the hexamer. We then docked the MTase domain to this complex and observed a compelling binding mode in which the MTase rotated approximately 180° with respect to its x-ray structure conformation (Figure 3b). The interface with nsp15 had excellent shape complementarity and established as many as five salt bridges, six hydrogen bonds, and a cation-pi interaction. The C-term zinc finger interacted with another antiparallel dsRNA as it passed over an EndoN site (on A1), while the C-term helix (residues 515-526) sat in the major groove of this same dsRNA. This set up a situation where three zinc fingers (two from nsp10 associated with another subunit and one from the nsp14 CTD) converged around the dsRNA just past the EndoN site. From this starting point, the nsp14/nsp15 structure was completed by building back in the 14-residue loop connecting the two nsp14 domains. Once fully optimized, the six nsp14/nsp10 subunits formed trimeric rings around each face of the nsp15 hexamer.

**Figure 3.**
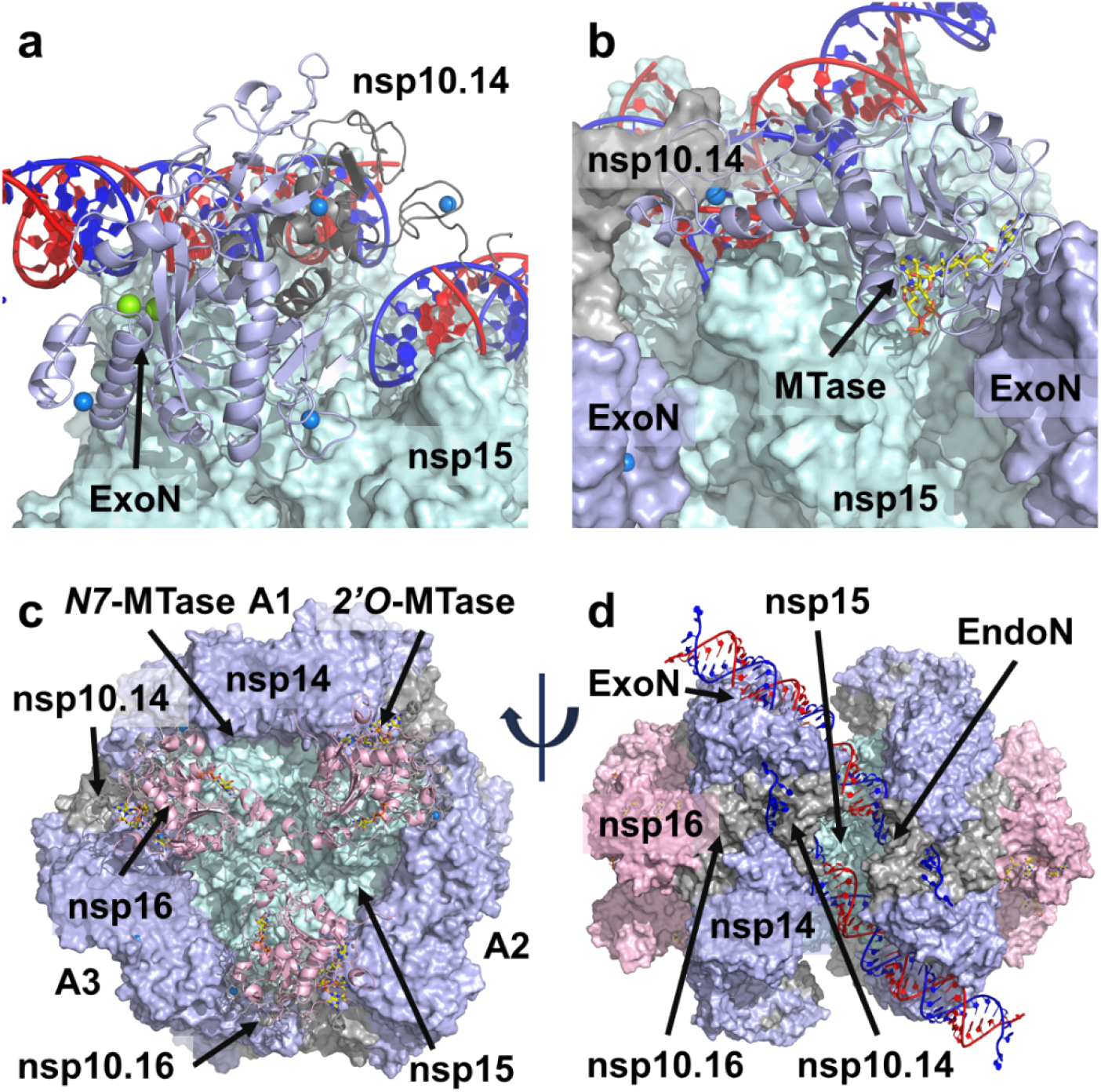
Formation of the nsp15/nsp14/nsp16/nsp10 complex. (a) The nsp14 ExoN NTD and nsp10 were manually positioned to interact with the dsRNA on the nsp15 hexamer, following the observed Lassa ExoN interaction with dsRNA. Nsp10 is positioned such that its two zinc fingers are over an antiparallel dsRNA, just past the nsp15 EndoN site. Six ExoN/nsp10 subunits can be arranged around the nsp15 hexamer. (b) The nsp14 MTase CTD was docked to the nsp15/ExoN/nsp10 hexameric complex. The CTD zinc finger is positioned over an antiparallel dsRNA, opposite the nsp10 associated with another nsp14 subunit. The binding mode reflects a significant conformational change between the two domains of nsp14. (c) Nsp16/nsp10 is docked to the nsp15/nsp14/nsp10 hexameric complex. Nsp10 is positioned between two nsp14 subunits, while three nsp16 subunits meet in the middle of the nsp15 trigonal face. (d) The full nsp15/nsp14/nsp16/(nsp10)_2_ hexamer.

### Nsp16/nsp10-nsp14-nsp15 interaction

After optimizing the nsp15 and nsp14/nsp10.14 interaction, we used protein-protein docking to investigate possible binding locations for nsp16/nsp10.16. From this, we identified an unambiguous site in which nsp10.16 was positioned between the nsp14 ExoN domain of A1 and the MTase domain of A2 (Figure 3c). Three such binding sites exist on each face of the hexamer, leading to the three nsp16 subunits lightly contacting in the center of each face. Key interactions included as many as six salt bridges and coordination of nsp16 D102 to the nsp14 ExoN zinc finger at C206, C209 and C225. As docking was performed without the nsp10.16 N-term residues 1-23 and C-term residues 132-139, these were subsequently built in during optimization. The flexible nsp10.16 N-term helix (residues 11-20) easily fit into a pocket formed by the nsp14 ExoN and its associated nsp10.14 subunit. With the placement of both nsp14/nsp10.14 and nsp16/nsp10.16, a ring of three nsp14/nsp16/(nsp10)_2_ subunits cap one face of the nsp15 hexamer, with an equivalent ring on the opposite face. Each of these rings establishes an interaction between its three ExoN domains and three dsRNA helices, while also positioning clusters of zinc fingers around the three antiparallel dsRNA helices just above the EndoN sites (Figure 3d).

The significance of this placement of nsp16/nsp10.16 with respect to nsp15/nsp14/nsp10.14 is that it puts both MTase sites near each other, an outcome consistent with their roles in capping. It also adds an additional pair of zinc fingers to interact with the dsRNA just past the EndoN site. In total there are six of these in close proximity (two from nsp10.14, two from nsp10.16, one from nsp14 ExoN, and one from nsp14 MTase). Interestingly, a narrow channel lined with basic residues is carved out by this arrangement of the proteins that could accommodate ssRNA. It runs from the EndoN site to the trigonal face accommodating both MTase sites. Given the dsRNA steric blockade also created by these proteins, we concluded this was the site of strand separation. Without the need for a helicase, the model suggests strand separation is facilitated by the zinc fingers, three of which (two from nsp10.14 and one from nsp14 MTase) act to direct the 3’ strand away from the complex, while two others (one from nsp10.16 and one from nsp14 ExoN) direct the 5’ strand into the basic channel. From there, the 5’ strand is funneled to the mRNA capping sites (nsp14 MTase and nsp16 MTase). We built a model of RNA following these two paths to illustrate the point.

### Nsp12/(nsp8)_2_-nsp14/nsp15 interaction

As discussed above, the most likely site for the ExoN and the polymerase to interact was on the exit face of nsp12, with the ExoN sitting below the dsRNA. This position of the ExoN relative to the polymerase active site has precedent in the *E. coli* DNA polymerase I Klenow fragment,[41] although the proposed binding orientation is not strictly identical. Alternative locations had been considered, based on other DNA polymerase structures which contain an ExoN domain, but these were ruled out by clear clashes with either nsp8 or nsp13. But as with nsp14/nsp15, docking of existing structures of nsp14 were unsuccessful in finding a suitable binding mode. The ExoN domain could be manually positioned to a satisfactory degree, but the MTase domain could not be properly placed.

This situation changed dramatically with the conformational change to nsp14, facilitated by binding to nsp15. This new conformation makes a significantly different surface available for binding to nsp12. With additional steric constraints coming from the nsp8 helices of the polymerase complex, which extend along the exiting dsRNA, a single binding mode between nsp12 and nsp14/nsp15 became the only choice and was further optimized. This binding position fuses the dsRNA of the nsp14/nsp15 complex with the dsRNA of the polymerase complex, placing the ExoN directly below the RNA as it exits nsp12 (Figure 4a). The MTase domain binds to the surface of nsp12 created by the helical C-term residues from 855-923, establishing multiple hydrogen bonds and salt bridge connections. The unusual beta hairpin (815-831) that protrudes from nsp12 below the dsRNA is positioned in the cleft between the two nsp14 domains (Figure 4b). Notably, D825 on this beta hairpin is in proximity to another nsp14 zinc finger, which sits below the ExoN active site. This complex was optimized, allowing additional flexibility for the poorly resolved nsp12 C-term residues 917-929.

**Figure 4.**
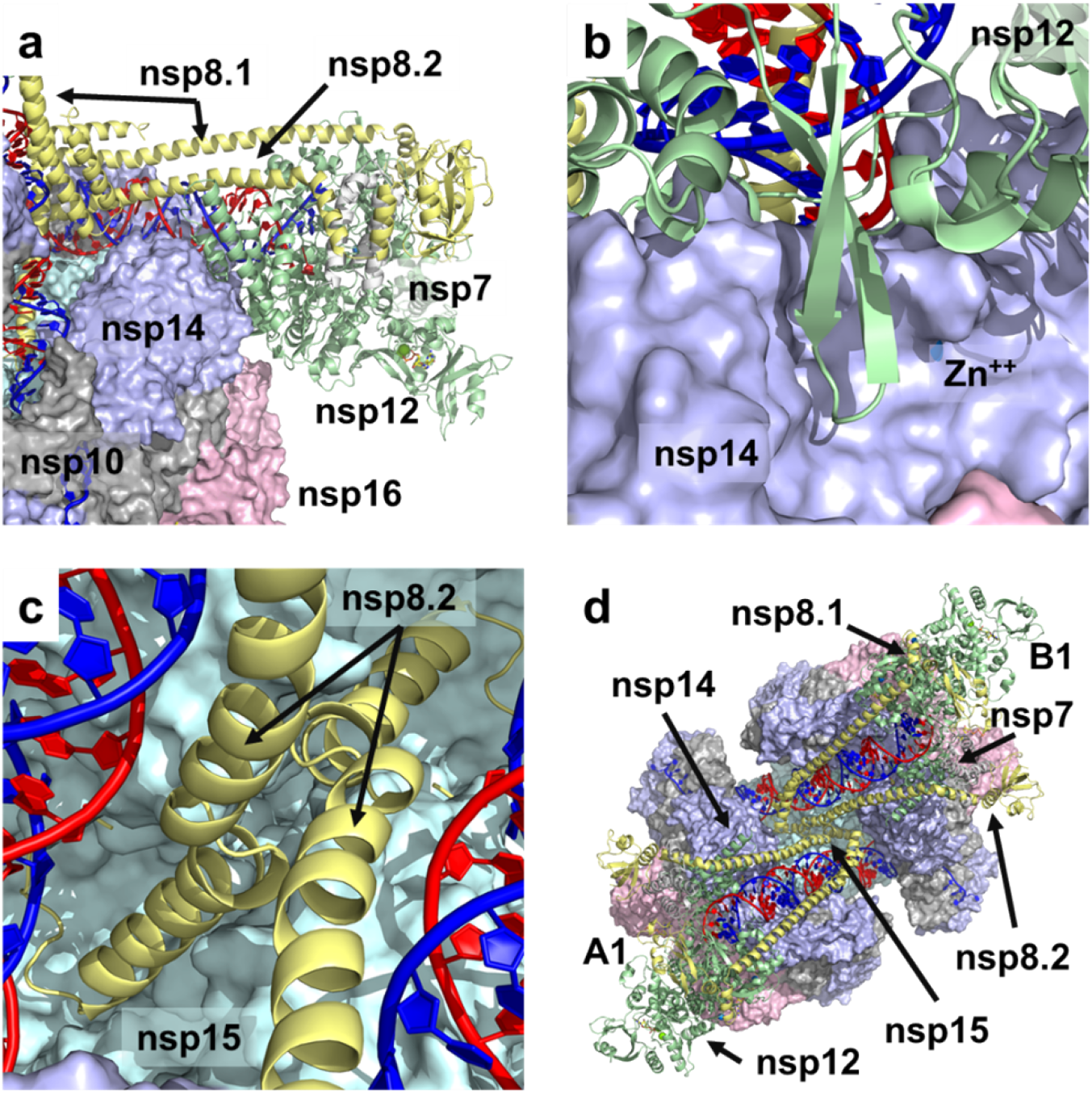
Binding of the polymerase to the nsp15/nsp14/nsp16 complex. (a) Binding of the polymerase is largely through nsp12 interactions with the conformationally altered face of nsp14. Much of this binding comes from the C-term helices of nsp12 (residues 855-923) interacting with the MTase domain of nsp14. (b) The nsp12 beta hairpin (residues 815-831) sits in the cleft between the two domains of nsp14, in close proximity to a zinc finger. (c) The short N-term helix of nsp8.2 (residues 12-28) extends to interact with nsp15, with the N-term residues (1-11) sitting under the dsRNA just ahead of the EndoN site. (d) View of a pair of polymerases bound to the nsp15/nsp14/nsp16 complex. One polymerase complex is associated with the A1 nsp14 subunit, while the other is associated with the B1 subunit. Their nsp8.2 subunits meet in the middle where they interact with nsp15.

When arranging additional polymerase complexes around the hexamer, we then focused on the position of the nsp8 N-term helices which extend over the dsRNA. These helices are composed of a short helix (residues 12-28) and a long helix (residues 32-97), which are seen to fold back into a bundle via a short connecting loop (residues 29-31). The helices of nsp8.1 did not appear to provide any new contacts with other proteins of the complex, stopping just short of touching nsp14 on the opposite face. However, nsp8.2 appeared to have a close contact with nsp8.2 coming from the opposite face (e.g. subunits A1 and B1). This interaction occurs in the center of the complex at the loop connecting the long and short nsp8 helices (residues 29-31). We initially considered optimizing the structure in this state, however, given direct binding of MERS nsp8 to nsp15 has been observed,[27] as has cellular colocalization of SARS nsp8 and nsp15[39], we considered the potential for a greater interaction between nsp8.2 and nsp15, with the most likely point of contact coming from the short helix and N-term residues. We thus docked the short nsp8 N-term helix to the nsp15 hexamer and identified a site in the center of the complex which would be consistent with a nearly linear extension of the short nsp8 helix from the longer nsp8 helix. Optimization of this extended state positioned the N-term residues on nsp8.2 under the dsRNA just prior to it passing over the EndoN site (Figure 4c). This set up a situation where the two nsp8.2 helices coming from opposite faces crossed in the center of the complex. Optimization of this crossing was facilitated by symmetric interactions between nsp15 R138 and nsp8 D30 and E32 on each subunit. This structure established a framework of pairs of polymerase complexes binding through nsp14 to the hexameric nsp15 complex (Figure 4d). Three such pairs can be arranged around the hexamer.

### Arrangement of nsp13

The position of nsp13 in the complex is already established by multiple cryo-EM structures which show that two nsp13 subunits can coordinate to each nsp12 polymerase through the nsp8 subunits.[5, 32, 36, 45] However, it is unclear if that will occur in the larger complex. Polymerases with a single nsp13 subunit associated, as well as structures with no associated nsp13, have been observed.[32] Considering nsp12, nsp13, nsp14, nsp15 and nsp16 stem from a single polyprotein and are expected to be generated in equal quantities, it is likely that six nsp13 subunits associate with the hexameric RTC. This is reinforced by the pairwise arrangement of nsp12 subunits on the complex. The proximity of these subunits, in and of itself, does not lead to clashes, such that four nsp13 subunits could theoretically be accommodated on a single pair of polymerases (twelve in total on the complex). However, as discussed below, we expect that the N-protein bound nsp13 dimer will coordinate the bulky 5’-UTR RNA, which appears unlikely for a complex saturated with nsp13 subunits. If the number of nsp13 subunits is limited to two for each polymerase pair, no such spatial limitations exist. Thus, the two nsp13 subunits can be arranged across polymerase pairs either by situating both on a single polymerase or distributing one on each. We propose that six nsp13 subunits are distributed among the three polymerase pairs as follows: two on A1 and none on B1; one on A2 and one on B2; and none on A3 and two on B3 (Figure 5). The two polymerases that coordinate two nsp13 subunits each (A1 and B3) bind the N protein, positioning the TRS-L above the Pol active site. These two nsp12 subunits would thus be in a configuration appropriate for transcription. The two polymerases that coordinate only a single nsp13 subunit (A2 and B2) would only be capable of replication. Thus, this arrangement would support four simultaneous polymerization events: two driving negative-strand synthesis with template switching (transcription), and two driving positive-strand replication.

**Figure 5.**
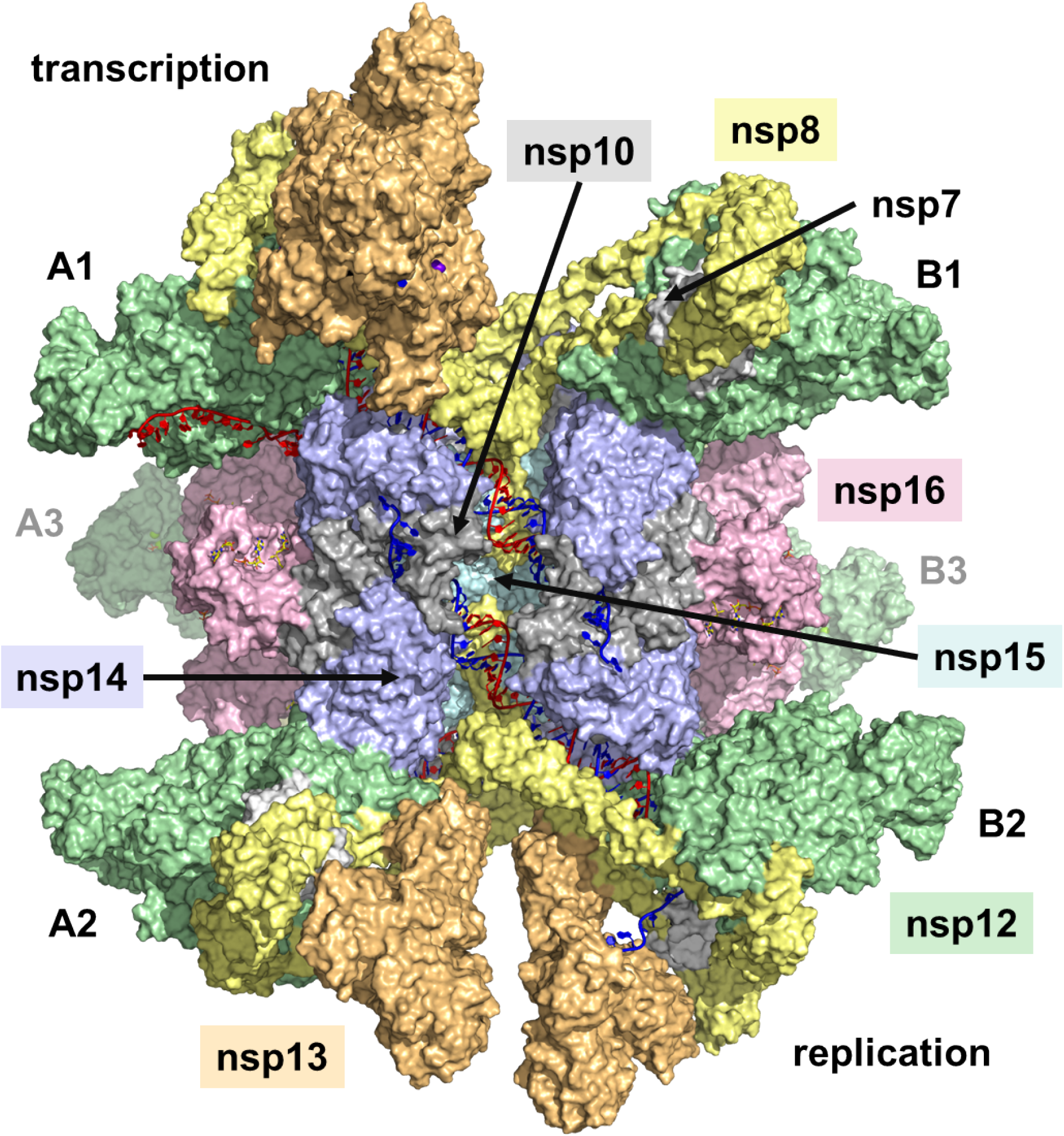
The complete replication/transcription complex, with a stoichiometry of 6 nsp15, 6 nsp14, 6 nsp16, 6 nsp12, 6 nsp13, 6 nsp7, 12 nsp8, 12 nsp10 and 2 N proteins. The six nsp13 subunits are arranged across nsp12 pairs in 2/0 (A1/B1), 1/1 (A2/B2), and 0/2 (A3/B3) stoichiometries. The polymerase complexes with two associated nsp13 subunits (A1 and B3) bind the N protein with the TRS-L oligo and are responsible for template switching during negative-strand synthesis (transcription). The two polymerase complexes with a single nsp13 subunit (A2 and B2) are responsible for replication.

## Discussion

### Overall organization

The structure of the SARS-CoV-2 RTC as outlined above centers around the nsp15 hexamer, which is often viewed as non-essential, because inactivating mutations within its EndoN active site are not lethal.[38] However, this can be deceptive, as the protein could still provide an important structural component beyond its enzymatic activity. Indeed, these same studies demonstrated that a mutation beyond the EndoN active site is in fact lethal to the virus. This D324A mutation found in murine hepatitis virus (MHV) corresponds to D296 in SARS-CoV-2, and likely leads to misfolding of the protein, causing it to be insoluble.

We find that three subunits each of nsp14/nsp10 and nsp16/nsp10 combine to form trigonal caps, two of which bind to opposite faces of the nsp15 hexamer. This leads to an overall nsp15:nsp14:nsp16:nsp10 stoichiometry of 6:6:6:12. To achieve this, nsp14 undergoes a significant conformational change where the MTase domain rotates approximately 180° relative to its x-ray structure conformation. While a small degree of conformational flexibility of this protein has been observed in the available SARS-CoV x-ray crystal structures, a much larger conformational change has recently been demonstrated via MD simulations.[17] The conformational change we observe here is energetically unfavorable for isolated nsp14, but is compensated by binding to nsp15 and nsp16.

The conformational change that nsp14 undergoes upon complexing with nsp15 and nsp16 enables its binding to nsp12. Much of the nsp12-nsp14 interaction occurs on the newly exposed face of the nsp14 MTase domain and within the cleft between the MTase and ExoN domains. The ExoN domain is positioned to interact with the newly formed dsRNA as it exits the polymerase, providing the foundation for proofreading. The arrangement of the polymerase subunits around the complex can be viewed as three sets of pairs. Within each pair, the nsp8.1 N-term helices are seen to cross in the center of the complex, extending to gain an additional interaction with nsp15.

We conclude that the likely association of nsp13 is a distribution of six subunits around the hexameric complex, retaining the expected molar ratios from the *ORF1ab* polyprotein. These can be arranged either with two nsp13 subunits on one nsp12 subunit and none on the partnered nsp12 subunit, or with one each on both nsp12 subunits. As we showed that the N protein NTD binds to the polymerase complex between two nsp13 subunits, positioning the TRS-L RNA above the polymerase active site, this situation would be suitable for negative-strand synthesis, which involves template switching at the TRS. With no such requirement for positive-strand synthesis, the polymerase complexes with a single nsp13 subunit would be suitable for full genome replication. Thus, we propose an arrangement of the nsp13 subunits across the six polymerase structures in which two nsp12 subunits coordinate a pair of nsp13 subunits each, two nsp12 subunits coordinate a single nsp13 subunit each, and the other two nsp12 subunits coordinate no nsp13. Such an arrangement would accommodate four simultaneous polymerization events, two for positive-strand synthesis and two for negative-strand synthesis.

The resulting structure provides a detailed atomic level view of coronavirus RNA replication and transcription. It suggests an efficient process in handling the viral RNA. As depicted schematically in Figure 6a, upon exiting the Pol active site, dsRNA passes over the ExoN site of nsp14 and then the EndoN site of nsp15. At this point, it encounters a collection of proteins which includes nsp14, nsp10.14 and nsp10.16. The close proximity of several zinc fingers leads to strand separation, where the template strand is directed away from the complex, while the nascent strand is funneled to sites responsible for mRNA capping. We describe below the implications of the structure on the major known functions of the RTC, including proofreading, mRNA capping and template switching.

**Figure 6.**
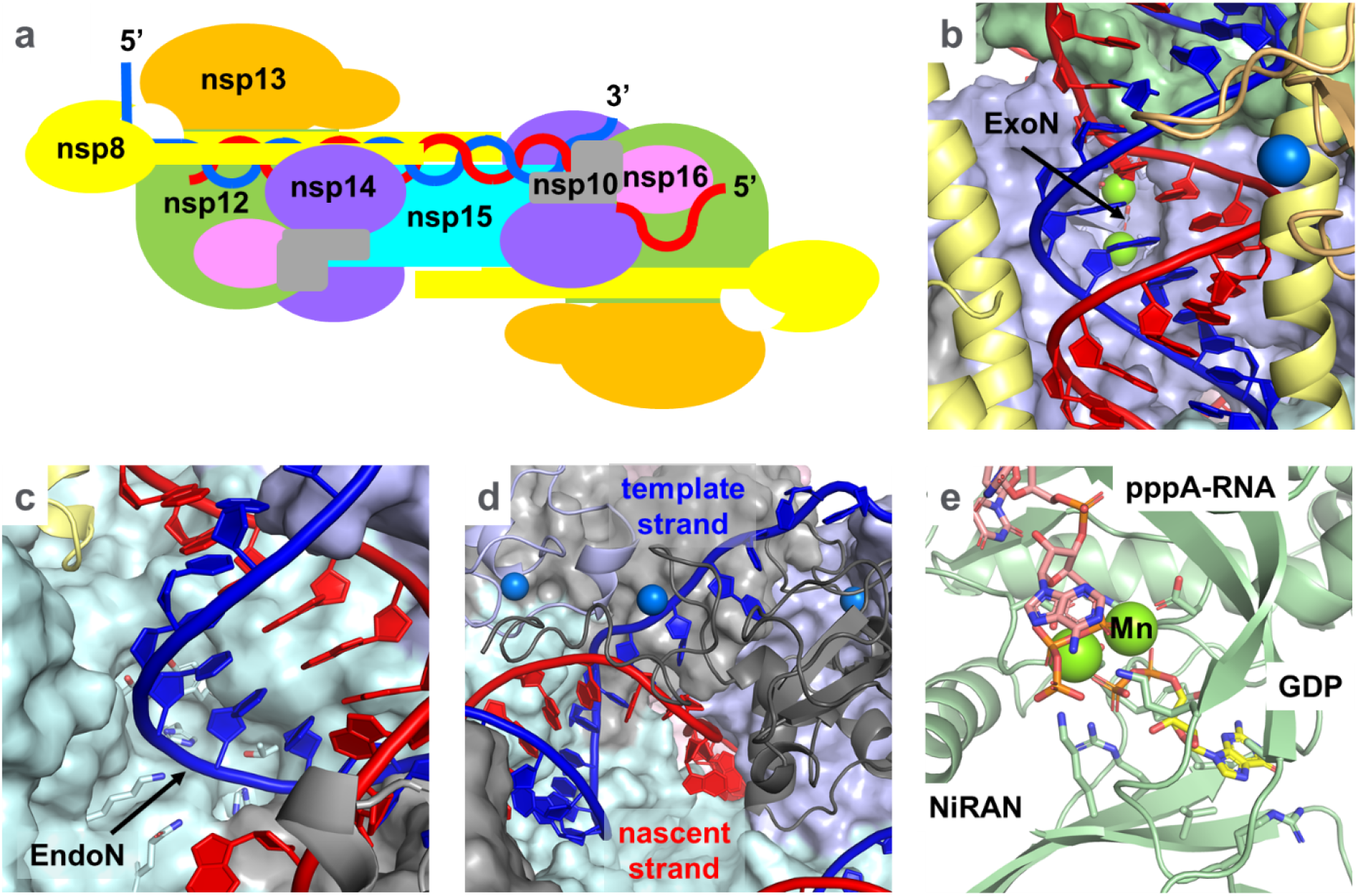
Details of some key functions. (a) Schematic representation of the RNA path. dsRNA makes its way from the nsp12 polymerase, across the nsp14 ExoN and nsp15 EndoN. It is separated into template (blue) and nascent (red) strands at nsp10, and the nascent strand is directed to the NiRAN and two MTase sites. (b) Detail of dsRNA exiting the polymerase and passing over the ExoN. The dsRNA is expected to shift into the ExoN active site when encountering a prematurely terminated nascent strand. (c) Detail of the dsRNA passing over the EndoN site, where the template strand would be the substrate. (d)Detail of strand separation occurring at the convergence of two zinc fingers from nsp10.14 and one from nsp14 CTD. The template strand is directed away from the complex, while the nascent strand is funneled down to the capping sites. (e) Detail of the NiRAN site. The first capping step occurs when the NiRAN site transfers GDP to the 5’ pppA-RNA, releasing a pyrophosphate.

### Backtracking

The structural picture of polymerization has already been well established. However, an important detail which still needs clarification is the exact role of nsp13. Its position above the polymerase active site is not consistent with dsRNA unwinding or a direct role in mRNA capping. Structurally, as discussed below, it is critical to coordination of the N protein needed for transcription. But its role as a traditional helicase appears questionable in the context of this model.

Cryo-EM structures have shown that the 5’ template overhang sits in the RNA binding groove of the helicase. But nsp13 is an SF1B helicase, with a polarity (5’ → 3’) that runs opposite to that of the polymerase (3’ → 5’), setting up a tug of war between the two. A solution to this conundrum has been suggested by a recent structure from Darst, et al.[46] which reveals nsp13 can adopt what appears to be an inactive state. In this state, the 1B domain undergoes a large conformational shift, opening up the RNA binding groove. This state is similar to the inactive state observed in the SF1B Pif1 DNA helicase from *Bacteroides spp*, where a rotation of the 1B domain was demonstrated to regulate activity of the helicase.[47] In its inactive form, polymerization can proceed unimpeded. However, when the protein is activated, the template would be expected to reverse course, leading to backtracking, a process important to both proofreading and transcription, as described below.

We should note that so far, the template has only been seen engaging with nsp13.1. However, binding of the N protein to the nsp13 dimer appears to block access to the nsp13.1 RNA binding groove. Modeling of the RNA template overhang into the nsp13.2 binding groove suggests this helicase should be capable of performing the same backtracking function. While not yet observed, the inactive state appears to be accessible to nsp13.2 as well. Thus, during positive-strand synthesis, the single nsp13.1 subunit is expected to control backtracking, while in negative-strand synthesis with bound N protein, nsp13.2 is expected to assume that role.

### Proofreading

Immediately upon exit from the polymerase, the newly formed dsRNA encounters the nsp14 ExoN (Figure 6b). This site is one complete helical turn from the polymerase active site. In an actively elongating state, the dsRNA is not expected to engage with the ExoN catalytic Mg^++^ ions, as it is prevented from doing so by steric interactions between the nascent strand and nsp14 P141 and G250 on loops that flank the active site. Examination of the homologous Lassa ExoN structure with bound dsRNA[44] indicates dsRNA can engage the ExoN site only at the 3’ terminal nucleotide. This 3’ terminus removes the steric constraints coming from the nascent strand, allowing the RNA to collapse into the active site. This suggests a scenario for proofreading in which a pause in nucleotide incorporation following a mismatched incorporation event would be subject to translocation of the prematurely terminated RNA to the ExoN site. The 3’ terminated RNA, now free of steric constraints, shifts into the active site to cleave the 3’ mismatched base.

Once the 3’ terminal nucleotide is removed, the RNA would need to be reset to continue polymerization. While it has been suggested that nsp13 driven backtracking may serve to initiate proofreading,[45] here we propose that backtracking serves to return the RNA to the polymerizing state. How backtracking would be triggered, is unknown, but a shift in the position of the RNA as it engages the ExoN may be the likely culprit. The coordinating nsp8 helices have proven themselves to be highly flexible and may serve to transmit changes in the RNA position to the nsp13 zinc binding domains (ZBD’s). When the RNA is out of position, nsp13 is activated. When the RNA is returned to the polymerase active site through backtracking, nsp13 resumes its resting state and RNA synthesis continues.

### Endonuclease activity

The nsp15 endonuclease is less well characterized than some of the other viral enzymes. It shows a preference for cleavage at U,[48] and dsRNA is a better substrate than ssRNA. Recent work by Baker, et al.[9, 49, 50] suggests the EndoN acts on the 5’ poly-U tail of the negative-strand to avoid host immune responses. Indeed, while positive-strand 3’ poly-A tails can reach lengths of 100-130 nt’s,[51] negative-strand 5’ poly-U tails are significantly shorter (9-26 nt’s). As with the nsp14 ExoN site, the dsRNA which crosses nsp15 from one face to the other appears to be sterically constrained from engaging the EndoN site (Figure 6c). The template RNA is positioned directly above the EndoN site and appears more likely to be the substrate than the nascent strand, but it is held aloft by basic residues across the length of the complex. A scenario in which the EndoN site might be engaged would be once synthesis is complete and the blunt end of the dsRNA has travelled far enough across nsp15 that it would no longer be supported by the full complement of basic residues. A shifting of the remainder of the dsRNA would occur, allowing it to engage the EndoN active site. The model suggests this would be ∼10 nt’s from the 5’ end of the template. This would be consistent with observations that nsp15 acts on the 5’ poly U tail of the negative-strand during positive-strand synthesis. Thus, once positive-strand synthesis is complete, the negative-strand poly U tail would be truncated.

### Strand separation

The proteins of the RTC feature multiple zinc fingers, whose role remains unclear. In total, there are twelve zinc fingers for each unit of the hexamer: three on nsp13, three on nsp14, two on nsp12 and two on each of the nsp10 subunits. Of these, two may be important for protein-protein binding: the nsp14 Zn^++^ at C206/C209/C225/H228 coordinates to nsp16 D102, and the nsp14 Zn^++^ at H256/C260/H263/C278 coordinates to nsp12 D825. But within the complex, we find a particularly interesting set of six of these zinc fingers, coming from four different subunits that are positioned in close proximity to the EndoN active site. Two of these come from nsp10.14 and two from nsp10.16, while another two come from two distinct subunits of nsp14: the Zn^++^ at C206/C209/C225/H228 in the ExoN domain and the Zn^++^ at H486/C451/C476/C483 in the MTase domain. As described below, the role of this cluster of zinc fingers becomes clear, considering the trajectory of the dsRNA as it crosses nsp15.

Nsp14 and nsp10.16 form a barrier which prevents the dsRNA from advancing beyond the EndoN site. But this area, rich in both the zinc fingers and basic residues, creates well defined pathways to separate the template and nascent strands (Figure 6d). The nsp14 C-term residues 519-526 sit in the major groove of the dsRNA, just above the EndoN site. Just beyond this point, the nsp14 CTD zinc finger and one of the nsp14.10 zinc fingers sit above the nascent strand, distorting its path. The template strand is sterically prevented from continuing in its dsRNA form and is directed between the two nsp10.14 zinc fingers away from the complex. The nascent strand is then directed into a channel lined with basic residues primarily coming from nsp14 ExoN. It encounters two additional zinc fingers (the nsp14 ExoN zinc finger and one of the zinc fingers from nsp10.16). This channel funnels the nascent strand into the region where the capping sites (the nsp12 NiRAN site and the two MTases) are found. This model suggests the nsp13 helicase is not involved in strand separation. It is more consistent with strand separation that occurs in negative sense viruses such as influenza.[52]

### mRNA capping

The capping of the 5’ end of the positive sense RNA occurs through a series of steps that involves transfer of a G, followed by two methylation events (pppN-RNA → GpppN-RNA → m^7^GpppN-RNA → m^7^GpppNm-RNA)[53, 54]. Methylation is facilitated by the nsp14 and nsp16 MTases, but the first step is less clear. There is some evidence that both nsp13 and the NiRAN site on nsp12 are involved.[3, 5, 8] Following conventional mRNA capping mechanisms, the general assumption is that nsp13 acts as a 5’ RNA triphosphatase (NTPase), converting pppN-RNA to ppN-RNA, and the NiRAN site acts a guanylyltransferase (GTase), transferring GTP to ppN-RNA and releasing pyrophosphate. However, while pppN-RNA has been shown to be dephosphorylated by the nsp13 NTPase site, it is a mediocre substrate, being completely inhibited by cellular level concentrations of ATP.[8] In our model, Nsp13 is also not in the general vicinity of the other enzymatic sites linked to capping, making it unlikely to interact directly with the RNA during capping. On the other hand, GTP is a good substrate for nsp13, suggesting there should be relatively high local concentrations of GDP.[8] Thus, we suggest pppN-RNA is not dephosphorylated prior to the first capping step, but instead the NiRAN site facilitates the transfer of GDP, releasing pyrophosphate, a mechanism more consistent with rhabdoviruses.[54]

Interestingly, the NiRAN site appears to have two functions. It is capable of nucleotidylating proteins with only a moderate preference for UMP,[4] and has also been shown to act on RNA.[5] Should it be responsible for the first step in capping, this would necessarily be G specific. While several cryo-EM structures of nsp12 have shown binding of nucleotides at the NiRAN site in a non-base specific manner, a recent cryo-EM structure with the guanosine analogue inhibitor, AT-527, identified a novel binding mode of the diphosphate form of the inhibitor in the NiRAN site that appears to specifically recognize its base.[55] With this as a starting point, we modeled the approach of pppA-RNA to the NiRAN site (Figure 6e).

After strand separation, the nascent strand is fed into the nsp15 trigonal face, where all three proposed capping sites are found (Figure S4). Charting its most likely initial pathway via conformational sampling, the RNA passes by three additional zinc fingers (a second nsp14 ExoN zinc finger and two on nsp12) before dropping down into the NiRAN active site. There, the pppN-RNA triphosphate binds to the same pair of catalytic Mn^++^ ions that bind to GDP. The orientation is such that a linkage between the β-PO_3_ of GDP and the α-PO_3_ of pppN-RNA would be formed upon release of pyrophosphate. It is unclear if this occurs in a single step or if a nucleotidylated protein intermediate is involved.

From this point the GpppN-RNA would continue to the nsp14 *N7*-MTase to become m^7^GpppN-RNA and then on to the nsp16 *2’-O*-MTase to become m^7^GpppNm-RNA (Figure S5). Further simulations are required to identify the probable path the RNA would take. There are several possible trajectories, as more than one site for each of these MTases is accessible and there are multiple areas across the entire capping region that are rich in basic residues to guide the RNA.

We note that it is not yet clear what distinguishes the positive-strand RNA from the negative-strand when it comes to the NiRAN site. It is possible they are not distinguished, and the negative-strand undergoes at least this first capping step, which would subsequently be cleaved through the EndoN, as described above. As an alternative, the model does not yet address the protein nucleotidylation capability of NiRAN and the role of nsp9. Given the negative-strand’s 5’ poly U tail and NiRAN’s preference for uridylation, attachment of the negative-strand to nsp9 should be considered as a mechanism to anchor the intermediate strand to the complex.

### Transcription

Discontinuous transcription is an unusual process in which a large segment of the template RNA is skipped over to create a set of nested subgenomic mRNA’s with a common 3’ end of varying lengths, all terminated with the same 5’ leader.[37] The shift in template is triggered when the TRS element is reached during negative-strand synthesis. As shown in Figure 1, this sequence of 7-10 nt’s is found several times throughout the genome: once in the 5’-UTR (referred to as TRS-L) and then preceding the starts of known ORF’s (referred to as TRS-B). When polymerization reaches the TRS-B, the newly synthesized complementary negative-strand recouples to the same template sequence in the TRS-L and completes synthesis of the 5’ leader, skipping over everything in between. The N protein has been thought to be involved in this process of recoupling, in that its NTD has been shown to specifically bind to the TRS sequence and is essential to transcription.[13, 56] The N protein CTD is a dimerization domain, such that binding of an N dimer to both the TRS-L and TRS-B would bring the two templates into close proximity. Yet a detailed picture of the mechanics of template switching is largely a mystery.

Here we showed via protein-protein docking that the N protein, when bound to the TRS oligo, positions itself between two nsp13 subunits over the polymerase active site. The orientation is such that once the complementary sequence is synthesized, the nascent strand could potentially recouple to this parallel template. We envision several factors that would allow this template switch to happen.

First, the N protein can bind to any TRS sequence (TRS-L or TRS-B), but specificity for TRS-L as the target of the template switch suggests additional *cis-*active RNA structural elements play a role. A series of papers on the SARS-CoV 5’-UTR structure established that SL2, which precedes the TRS-L, is required for transcription,[57] while SL4, which immediately follows the TRS-L, also appears to play a role.[58] Within the context of this model, these RNA structural elements likely provide additional binding to the nsp13 subunits, ensuring specificity for TRS-L over TRS-B. Preliminary modeling of SL2 to nsp13 confirms the plausibility of this hypothesis. Interestingly, positioning of the TRS-L over the polymerase active site may also aid initiation. Additional studies of the SARS-CoV 5’-UTR found SL1 interacts with the 3’-UTR, meaning the 5’ TRS-L should be in close proximity to the 3’ poly-A tail when it coordinates to the polymerase.[59] Thus, as with viruses such as influenza,[52] binding of the 5’-UTR may act as a promoter (Figure 7a).

**Figure 7.**
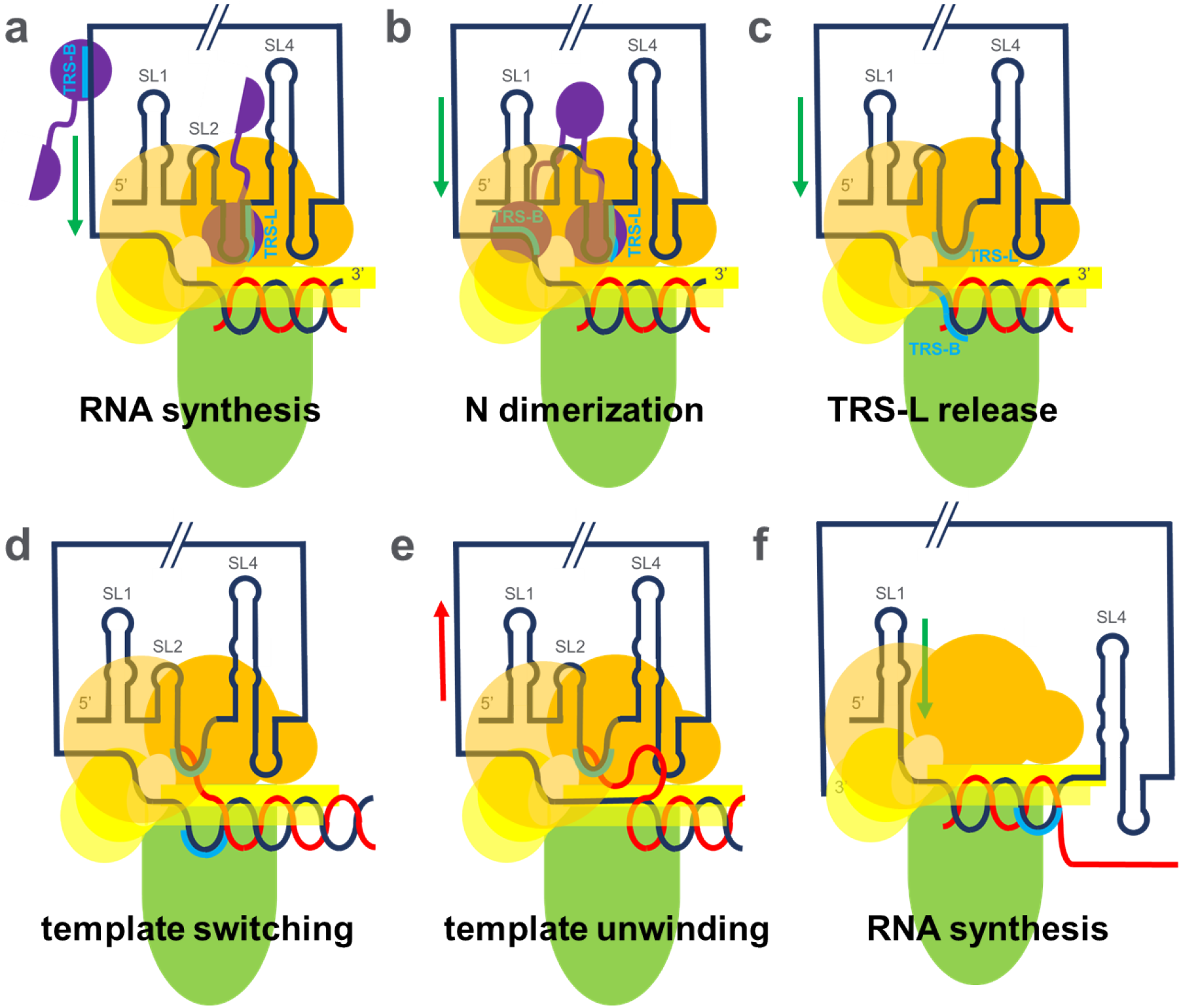
Model of template switching [nsp12 (green), nsp13 (orange), nsp8 (yellow), N protein (purple), template RNA (black), nascent RNA (red), TRS (blue)]. (a) The 5’-UTR coordinates to the nsp13 dimer, with TRS-L bound N protein positioned above the polymerase active site. RNA synthesis begins on the 3’ end of the template. (b) Synthesis continues until the N protein dimerizes with another N protein bound to TRS-B on the template. (c) The N proteins release the RNA. (d) The complementary TRS-B of the nascent strand recouples with TRS-L. (e) The shift in RNA position triggers nsp13 template backtracking, unwinding the dsRNA. (f) Once fully unwound, synthesis continues on the 5’ leader, starting from the TRS-L.

Second, the TRS-L is effectively sequestered by the N protein and would need to be released in order to base pair with the nascent strand. The N protein CTD is known to be a dimerization domain.[60] This raises the possibility that an N protein bound to a TRS-B would dimerize with the N protein bound to the TRS-L once that segment of the template reaches the polymerase (Figure 7b). This dimerization could then force the release of the TRS-L (Figure 7c), freeing it up for coupling to the nascent strand once the complementary sequence has been synthesized (Figure 7d).

Finally, the original template would need to unwind from the nascent strand for polymerization to continue on the new template. As with proofreading, we propose the shift in the dsRNA position upon base pairing of the nascent strand to the TRS-L triggers activation of nsp13 and backtracking of the template. This would unwind the original template from the nascent strand, while leaving it base paired with the new template (Figure 7e). Once unwinding is complete, the dsRNA formed at the TRS-L juncture with the new template could shift back into the polymerase active site and finish synthesis on the remaining nucleotides of the 5’ leader (Figure 7f).

### Summary

The detailed model of the SARS-CoV-2 RTC, which was derived in part from a series of protein-protein docking exercises, provides a comprehensive view of this complex and its efficient processing of the viral RNA. The model offers a framework for interpreting a range of observations, including proofreading, mRNA capping, and transcription. While the model is supported by a wide variety of experimental observations regarding coronavirus complex formation and function, we acknowledge it is only a starting point. More explicit simulations and experimental verification are needed to understand key details; models of other coronaviruses should be developed to confirm the primary structural interactions; and more careful consideration should be given to the roles of other viral proteins such as nsp9 as well as host factors. However, as it stands, we believe the model provides a new perspective on the complicated processes involved in coronavirus RNA replication, most importantly, serving as a guide for future assay development and structural studies.

## Methods

Structures used to construct the model were the following: nsp12/nsp7/nsp8/nsp13/dsRNA (PDB:6XEZ)[32]; nsp15 (PDB:6X1B)[25]; nsp16/nsp10 (PDB:6WVN) [21]; nsp9 (PDB:6W4B)[61]; and N NTD/TRS-L (PDB:7ACT)[31]. Since an x-ray structure of nsp14 was not available at the time this work was done, an homology model was built based on the SARS structure (PDB:5NFY, chains A/M)[6].

Protein-protein docking was carried out using the Piper[62] method within Bioluminate[63]. Refinement was an iterative process within the Schrödinger suite that involved sidechain and loop conformation optimization through Prime[64], minimization through Prime and Macromodel[65], and additional conformational sampling through Macromodel. Once the initial model was constructed, all proteins were replaced with their original PDB structures and reoptimized. The final structure was minimized in pieces, due to practical limits in dealing with such a large complex.

## Acknowledgments

The authors are grateful to Seth Darst for sharing structures in advance of publication and for many fruitful conversations which contributed to the development of this model.

## Supplementary Material

**Figure S1.**
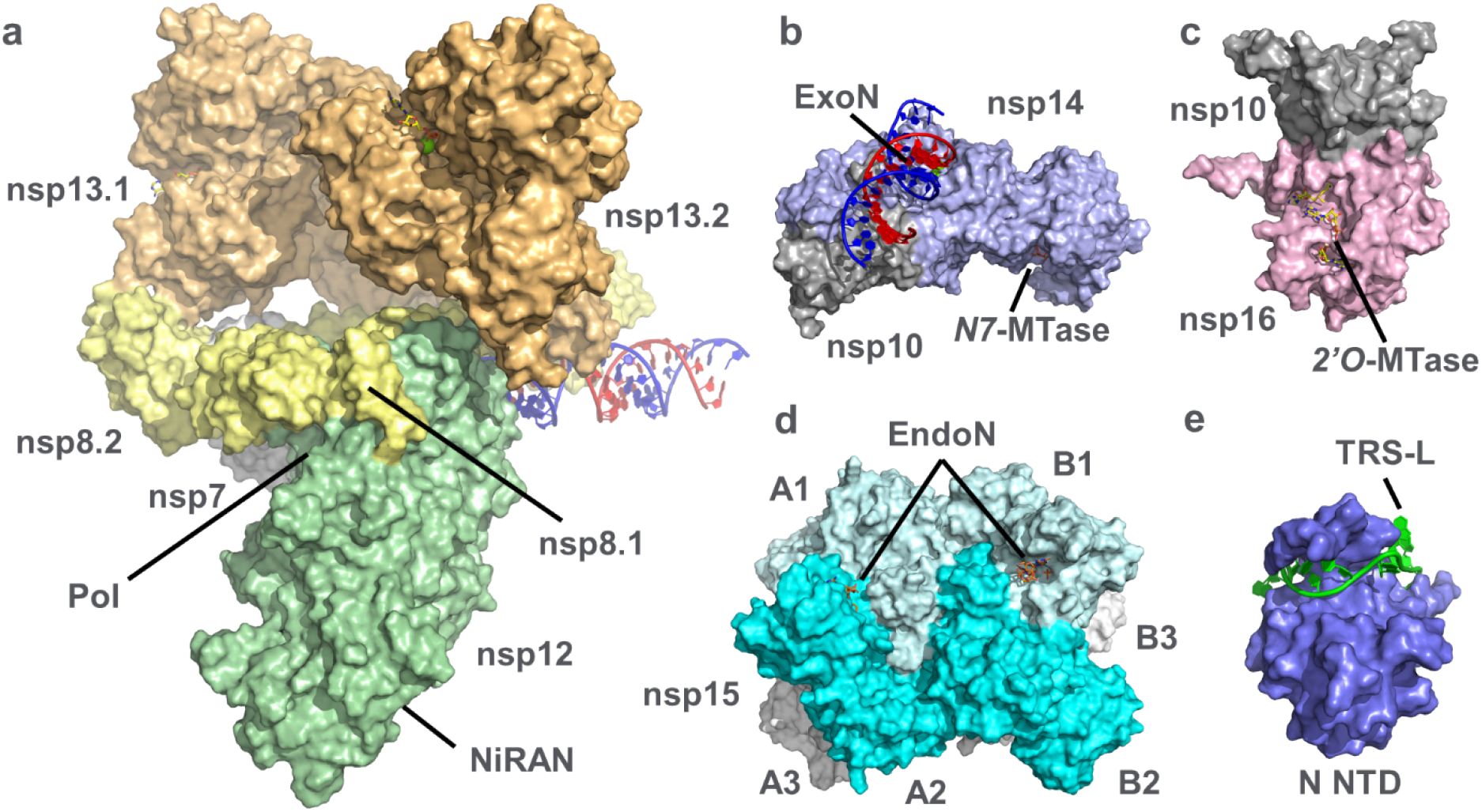
Key components of the coronavirus RTC. (a) SARS-CoV-2 polymerase complex of nsp12 (green), nsp13 (orange), nsp7 (white) and nsp8 (yellow) (PDB: 6XEZ). (b) Homology model of SARS-CoV-2 ExoN/*N7*-MTase nsp14 (blue) with nsp10 (gray) cofactor (based on SARS-CoV PDB: 5NFY, with dsRNA modeled after Lassa ExoN PDB: 4FVU). (c) SARS-CoV-2 *2’O*-MTase nsp16 (pink) with nsp10 cofactor (PDB: 6WVN). (d) Hexamer of SARS-CoV-2 EndoN nsp15 (cyan) (PDB: 6X1B). Subunits of the nsp15 hexamer are labeled, reflecting two trigonal faces (A1/A2/A3 and B1/B2/B3). (e) SARS-CoV-2 N protein NTP (residues 44-180) bound to the 10 nt TRS-L oligo (PDB: 7ACT).

**Figure S2.**
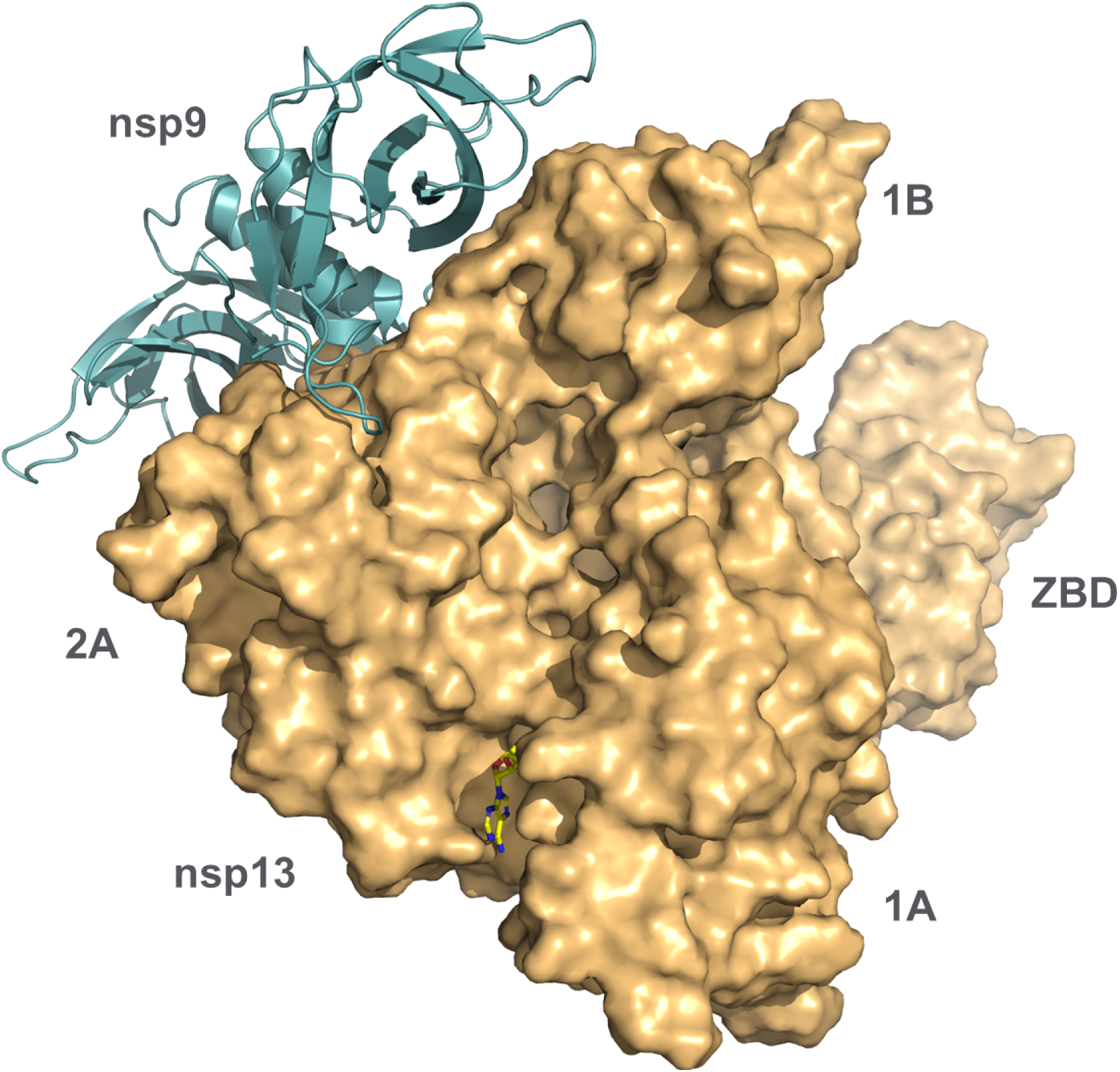
Docking of the nsp9 dimer to the active state of nsp13. Nsp9 coordinates to the 2A domain, where it is positioned to interact with the 5’ end of RNA as it exits the nsp13 RNA binding channel. A similar binding mode was found for other conformations of nsp13.

**Figure S3.**
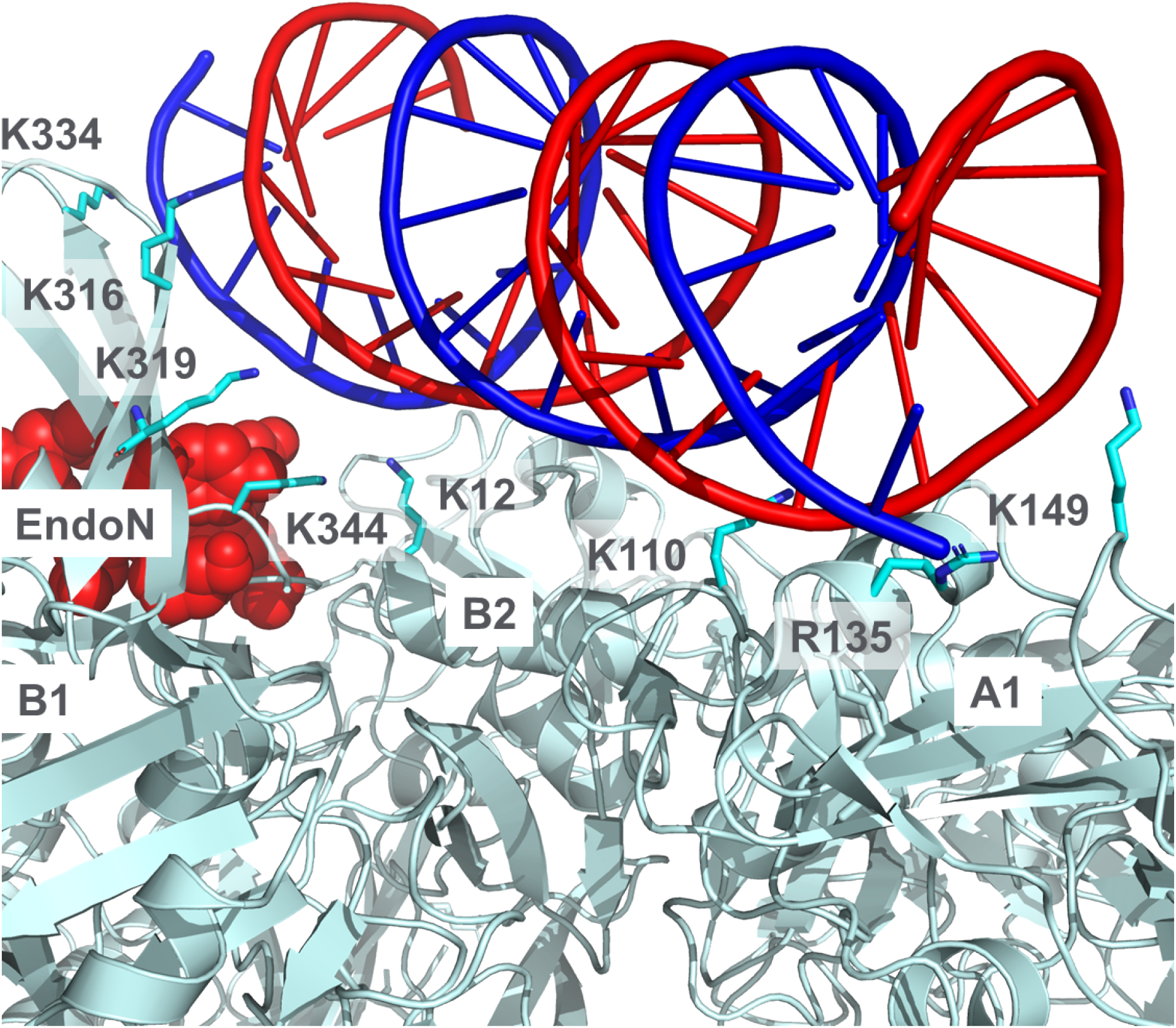
Detail of dsRNA path over hexameric nsp15. The RNA interacts with eight basic residues from three different subunits and passes over the EndoN site (red).

**Figure S4.**
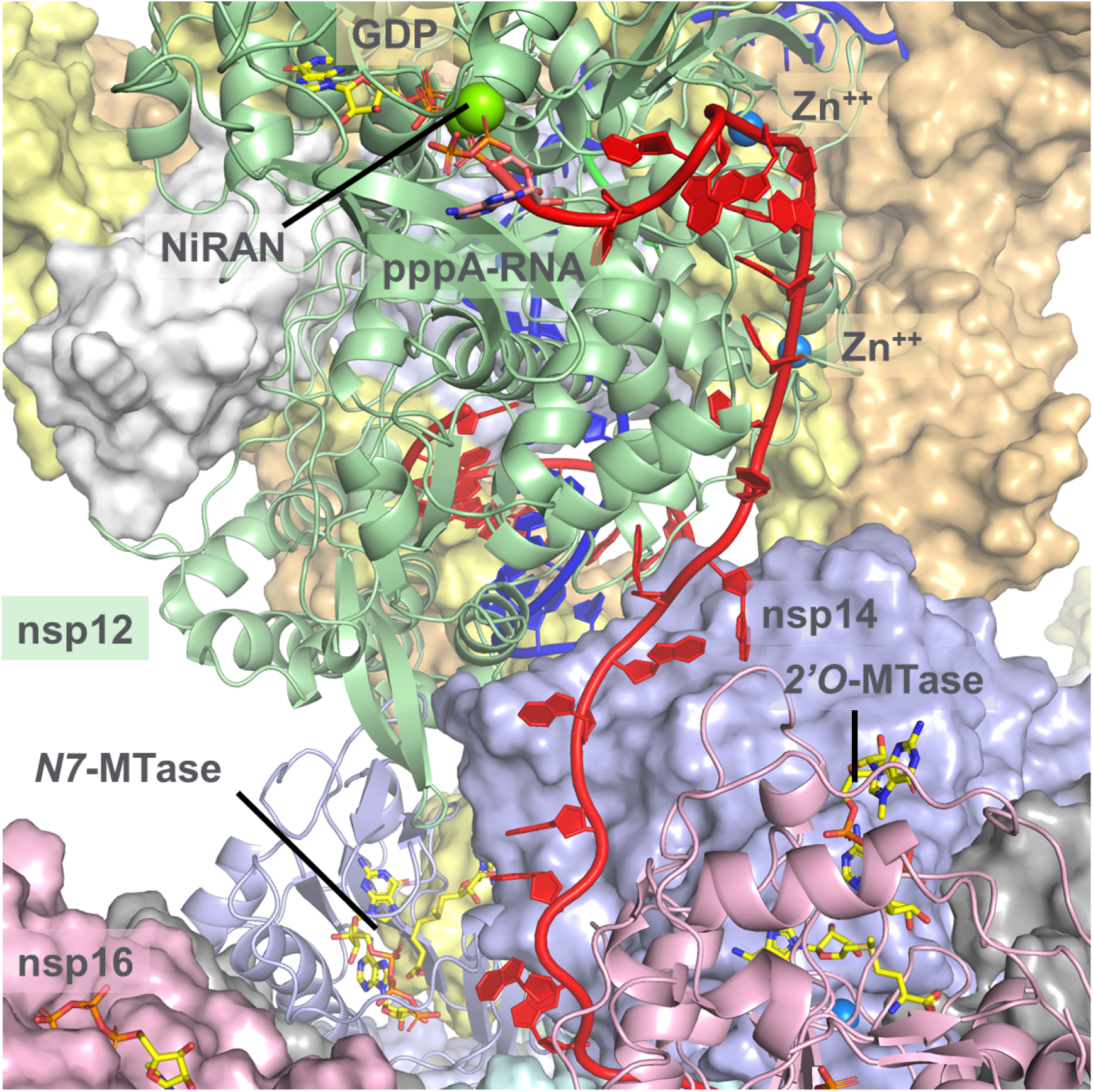
Detail of the nascent strand RNA emerging from a basic residue rich channel after strand separation and making its way to the NiRAN domain to initiate capping. The RNA passes by several zinc fingers, including two on nsp12. After the initial transfer of G to the pppA-RNA, the RNA makes its way to the nsp14 *N7*-MTase and nsp16 *2’O*-MTase to complete the mRNA capping.

**Figure S5.**
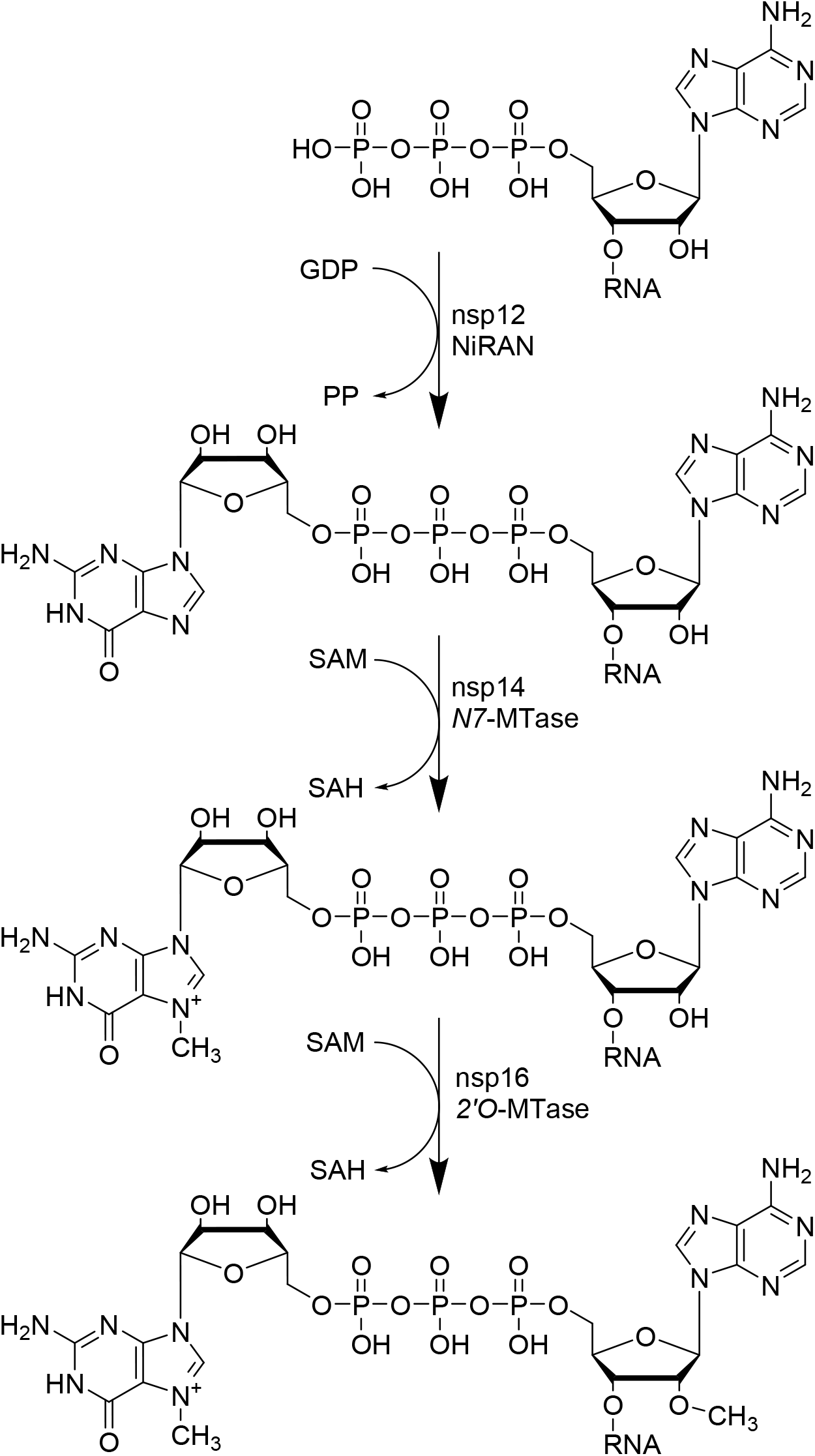
Proposed mRNA capping schematic. The nsp12 NiRAN site facilitates transfer of GDP to the 5’ pppA-RNA, with loss of pyrophosphate. The *N7* of the G is then methylated via the nsp14 MTase. Finally, the *O2’* of the A is methylated via the nsp16 MTase.

### PDB files

The hexameric RTC is divided into six separate PDB files.

Perry_SARS2-RTC_A1.pdb

Perry_SARS2-RTC_A2.pdb

Perry_SARS2-RTC_A3.pdb

Perry_SARS2-RTC_B1.pdb

Perry_SARS2-RTC_B2.pdb

Perry_SARS2-RTC_B3.pdb

### Movie file

3D view of the hexameric RTC. Perry_SARS2-RTC.mpg

